# Protein profiling reveals the characteristic changes of complement cascade pathway in the tissues of gastric signet ring cell carcinoma

**DOI:** 10.1101/816272

**Authors:** Yang Fan, Bin Bai, Yan Ren, Yanxia Liu, Fenli Zhou, Xiaomin Lou, Jin Zi, Guixue Hou, Qingchuan Zhao, Siqi Liu

## Abstract

Signet ring cell carcinoma (SRCC) is a histological subtype of gastric cancer that has distinct features in cellular morphology, epidemiology and clinicopathology compared with adenocarcinomas (ACs). Lacking of systematically molecular overview to this disease made a slow progress in diagnosis and therapy for SRCC. In the present proteomics study, the gastric tissues were collected from tumor and adjacent regions including 14 SRCC and 34 AC cases, and laser capture microdissection (LCM) was employed to eradicate cellular heterogeneity of the tissues. Over 6,000 proteins were quantified through data independent acquisition (DIA) mass spectrometry (MS). The quantitative profiles of proteomes in tumor tissues, either AC or SRCC, were dramatically different from that in the corresponding adjacencies, whereas the SRCC proteomes appeared not distinguishable to the AC proteomes via hierarchical clustering. However, focusing on univariate analysis and pathway enrichment unrevealed that some proteins and pathways bared the differences between SRCC and ACs. Importantly, the abundance changes for a bulk of proteins involved in complement cascade were highly associated with SRCC but not so sensitive to the AC status. A hypothesis, therefore, was proposed that the complement cascade was evoked in the SRCC microenvironment upon infiltration, while the SRCC cells survived from the complement cytotoxicity by secreting negative regulators. Moreover, an attempt was made to seek appropriate cell model for gastric SRCC, through proteomic comparison of the 15 gastric cell lines and the gastric tumors. The prediction upon supervised classifier suggested none of these gastric cell lines qualified in mimic to SRCC.

## Introduction

Gastric signet ring cell carcinoma (SRCC) is a histological subtype of gastric cancer defined by World Health Organization (WHO) as gastric tumors composed of predominantly or exclusively of signet-ring cells, which are characterized by a central optically clear, globoid droplet of cytoplasmic mucin with an eccentrically placed nucleus^1^. On the contrary to a trend of decreasing incidence of gastric cancer worldwide, the SRCC incidence has remained rising^2^. The molecular features of pathology and pharmacology relevant to SRCC are highly attractive in the frontier of gastric cancer study.

Gastric SRCC is not only special in its histology, but is also very different in clinicopathological features from other subtypes of gastric cancer. The female incidence of SRCC in all the gastric cancer is approximately 50%, whereas that of non-SRCC is about 30%; the average incidence age of SRCC is around 62 years, whereas that of non-SRCC is roughly 69 years^3^. Although *Helicobacter pylori* infection is regarded as a risk factor to gastric cancer, this bacterium is not commonly found in SRCC^4^. With comparison of SRCC to other two main subtypes of gastric cancer, well-moderately differentiated adenocarcinoma (WMDAC) and poorly differentiated adenocarcinoma (PDAC), Chon et al observed that at early stage the prognosis of SRCC was better than that of WMDAC and PDAC, whereas at later stage the SRCC prognosis was worse than other two subtypes^5^.

During the last decade, a number of studies dug the molecular indicators of gastric SRCC. Immunostaining revealed that all of gastric SRCC and mucinous adenocarcinoma with high abundance of trafficking kinesin protein 1^6^. The RT-PCR and IHC evidence demonstrated that the expression product of forkhead box P3 was significant upregulated in gastric cancer, especially the correspondent abundance with higher percentage in SRCC than in adenocarcinoma (79.3% versus 0%)^7^. Similar to the observations, several proteins such as pyruvate kinase M1/2, glypican-3, cathepsin E, and transmembrane protein 207 were found in abundance changes in the gastric SRCC cells or tissues. These studies touching the SRCC-related proteins are still at preliminary phase and are far from clinical practice. Most those proteins were individually divulged through different approaches and laboratories, and were not commonly verified. Proteomics as a powerful means in identification and quantification of proteins has naturally become a main technique for exploration of the SRCC-related proteins.

Proteomic investigation on gastric SRCC is still limited within a slow pace. There are only 3 published papers so far that discussed about SRCC using proteomics but did not reach any significant conclusion to help understanding of the molecular features of SRCC^8-10^. Which barrier did hinder the relevant studies to gastric SRCC? Three factors at least, according to our view, indeed affect the SRCC study. First of all, how to obtain a reasonable cohort of the SRCC samples is an obvious limit in this area. All the studies in the published literatures regarding the SRCC proteomics were only dealt with less than 4 cases and were less convincing for statistical evaluation. Secondly, how to excise the SRCC tissues is a key limit in the sample preparation. Since gastric SRCC owns its special histological features, the gastric tissues with dominant signet ring cells should be carefully estimated and isolated. Thirdly, how to conduct proteomic analysis is an important technique issue so that it provides a deep and large data in proteomic comparison, especially in a relatively large cohort.

With awareness of the 3 gaps, we presented in this communication, a comprehensive comparison of the proteomes derived from the gastric tissues of SRCC, PDAC and WMDAC. A cohort with 48 cases including 14 SRCC, 17 PDAC and 17 WMDAC cases was strictly selected from more than 2,500 cases and were carefully evaluated on the basis of histological examination. The cancer and adjacent tissues were well isolated using laser capture microdissection (LCM)^11^. We employed data independent acquisition (DIA)-based proteomics^12^ in quantitatively profiling proteomes for all the individuals at large scale. For the first time, the quantitative proteomes in gastric SRCC, PDAC and WMDAC were deeply characterized in parallel, revealing the proteins in the complement cascade pathway significantly upregulated in SRCC. Moreover, we made a proteomic survey to 14 gastric cancer cell lines aiming at classifying the cancer subtype-representativeness of each cell line.

## Methods

Detailed methods are presented in **Supplemental information 1**. In summary, frozen tissues of 3 subtypes of gastric cancer, SRCC, PDAC and WMDAC, were retrieved from Xijing Hospital, China, following the inclusion criteria described in **Figure 1**. Tumor cells and corresponding adjacent epithelial cells were isolated from tissue samples by LCM, while fifteen gastric cell lines were also collected from other laboratories and commercial sources (**Table S1**). The LCM samples and cell line samples were analyzed by DIA MS. The criteria in **Figure S2** were set to filter the data and identify tumor/adjacent differentially expressed proteins (T/A-DEPs). Statistical evaluation and protein fold changes were used to determine SRCC/AC DEPs (S/A-DEPs). Biologically relevant pathways were extracted by enrichment analysis. Machine learning based models were trained and used to predict tissue representativeness of gastric cell lines. All the MS data were deposited to the Chinese National GeneBank Sequence Archive (CNSA) database (https://db.cngb.org/cnsa/) (CNP0000652).

**Figure 1.**
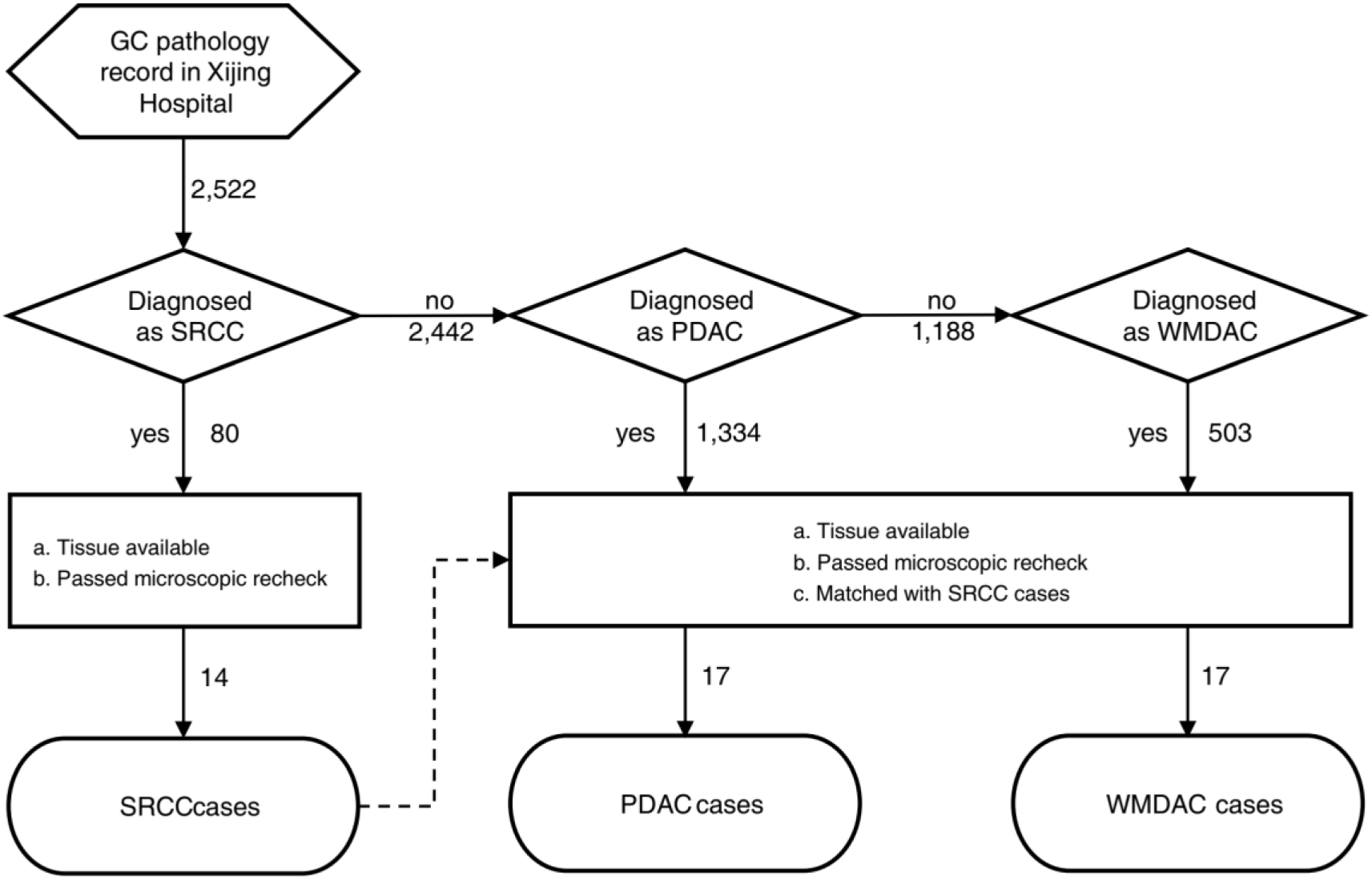
The evaluation procedure to select proper tissue samples of SRCC, PDAC and WMDAC for the proteomics study using LCM.

## Results

### Collection of high-quality cancer tissues and proteomic data

Of 80 SRCC cases recorded in the tissue bank of Xijing hospital, only 14 tumor tissues were qualified with major tumor cells with characters of signet ring type by H&E staining recheck (**Figure 1**). In the tissue bank, 2442 cases were primarily diagnosed as PDAC and WMDAC, 685 cases were removed after recheck, resulting in 1,334 PDAC and 503 WMDAC cases. For each subtype, 14 cases were selected in a case-wise matching manner regarding the 14 SRCC cases. Then 3 low-age cases with available frozen tissues and > 50% cancer cells for each subtype were supplemented, resulting in inclusion of 17 cases for PDAC and WMDAC. In total of 48 cases of gastric cancer that were well collected paired tissues of tumor and adjacent, these samples were histologically classified into SRCC (n = 14), PDAC (n =17) and WMDAC (n = 17). In order to set a base for cross-subtype comparison, the matching of clinicopathological features were specially considered in the selected cases. As a result, the age, gender, T and N staging of SRCC case were not significantly different from those of PDAC or WMDAC cases (**Table 1**). Of the 3 subtypes, the mean ages ranged from 54.79 to 58.35 years, the percentages of male cases ranged from 64% to 71%, the tumors were all in advanced stages, i.e. the T2, T3 and T4 stages, and the percentages of cases in their N0, N1/N2 and N3 stages ranged from 0% to 7%, 21% to 35% and 59% to 71%.

**Table 1.**
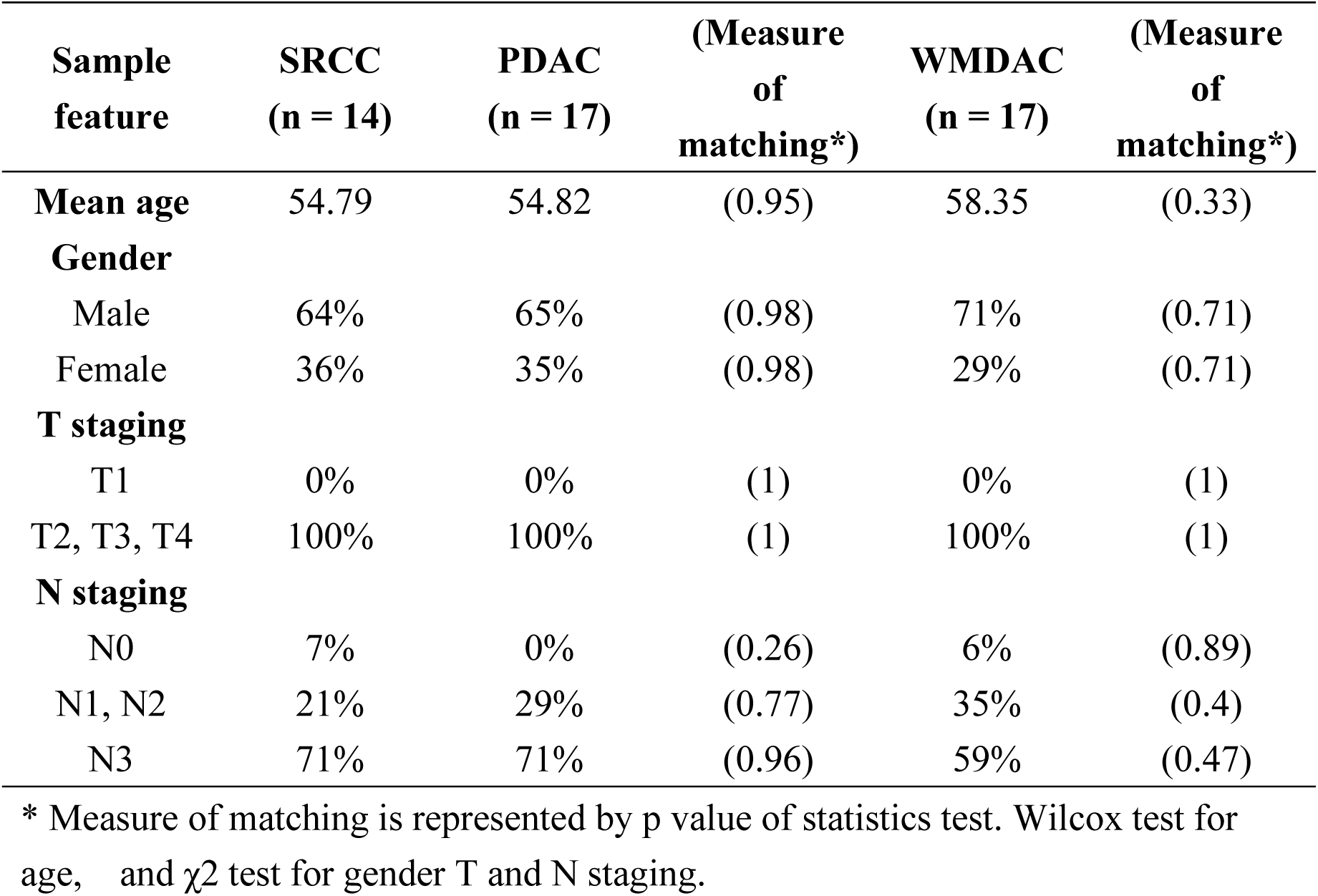
Statistics for tissue sample collection and pairing

| **Sample feature** | **SRCC**<br>**(n = 14)** | **PDAC**<br>**(n = 17)** | **(Measure**<br>**of**<br>**matching\*)** | **WMDAC**<br>**(n = 17)** | **(Measure**<br>**of**<br>**matching\*)** |
| --- | --- | --- | --- | --- | --- |
| **Mean age** | 54.79 | 54.82 | (0.95) | 58.35 | (0.33) |
| **Gender** |  |  |  |  |  |
| Male | 64% | 65% | (0.98) | 71% | (0.71) |
| Female | 36% | 35% | (0.98) | 29% | (0.71) |
| **T staging** |  |  |  |  |  |
| T1 | 0% | 0% | (1) | 0% | (1) |
| T2, T3, T4 | 100% | 100% | (1) | 100% | (1) |
| **N staging** |  |  |  |  |  |
| N0 | 7% | 0% | (0.26) | 6% | (0.89) |
| N1, N2 | 21% | 29% | (0.77) | 35% | (0.4) |
| N3 | 71% | 71% | (0.96) | 59% | (0.47) |

\* Measure of matching is represented by p value of statistics test. Wilcox test for age, and $\chi^2$ test for gender T and N staging.

Tumor cells are generally in uneven distribution in a resected tumor tissue. To obtain the tissues with high contents of tumor cells, we adopted LCM and collected the tissues with low intra-tumor heterogeneity for protein extraction. The typical microscopic images of the LCM treated tissues were presented in **Figure 2A**, 3 cases randomly selected from each subtype of gastric cancer, clearly demonstrating the “signet ring” morphology of SRCC, the densely formation of separate tumor cells of PDAC as well as the gland-like structures formed by WMDAC cells.

**Figure 2.**
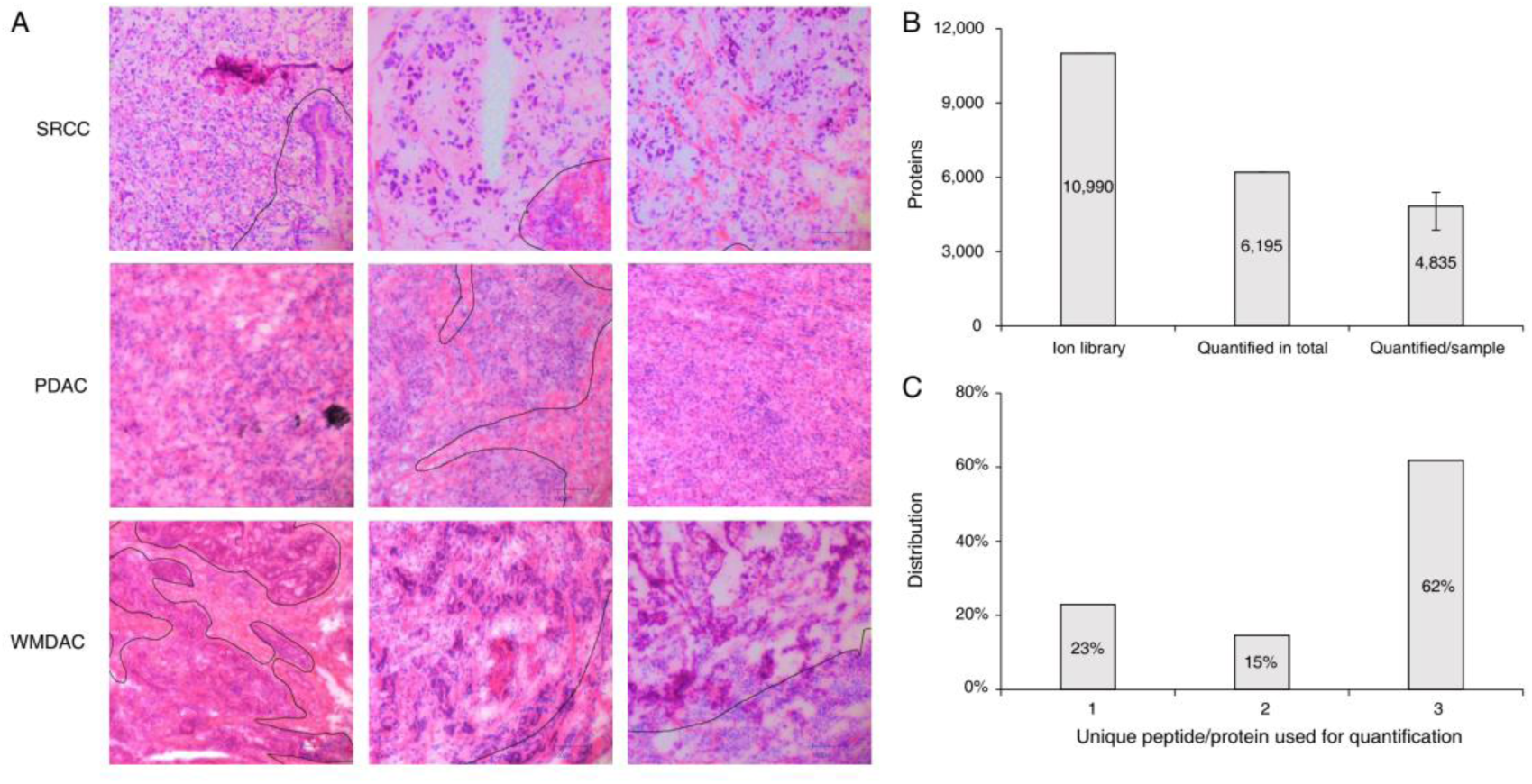
Assessment of data quality. A) The HE images for diagnosis of SRCC, PDAC and WMDAC. B) The proteins in the gastric tissues identified using DDA and DIA approach (error bar indicates the upper and lower bound of proteins quantified/sample). C) Distribution of the unique peptides in the quantified proteins.

The LCM samples with approximate area of 20 mm^2^ were processed through an established method in our laboratory that was suitable for extracting peptides from micro amount biological samples (**Methods**). A range of 1.4 to 12.5 μg peptides were retrieved from an LCM sample, and the peptide yields were ranged from 0.14 to 0.65 μg/mm^2^ LCM sample (**Table S3**).

For the sake of better protein identification, we employed Preview to rapidly interrogate the occurrences of 25 chemical modifications in the samples. The assessment results in **Table S2** surprisingly suggested that carbamidomethylation artifacts (+57 on N-terminus, H and K), deamidation (+1 on N and Q) and DTT addition (+152 on C) were top 3 modifications, while pyroglutamate formation (−17 on N-terminal Q) and oxidation (+16 on M), were ranked at the 6^th^ and 7^th^ places. Therefore, the top 3 modifications were set as variable modifications in subsequent database searching. The spectra library required by DIA analysis was constructed by merging the DDA search results from the samples that were treated with pooling and fractionation, and the publicly available pan human library^13^. The library covered 10,990 proteins, 163,254 peptides and 299,808 precursors, correspondingly.

For quality control of proteomic data, we checked for potential batch effect by feeding the unprocessed quantification data to principal component analysis (PCA). As visualized in **Figure S4**, no obvious deviation was found among the 4 sequential batches, which were gained from the continuous runs lasting a month. This implicated that the data quality was solid and batch effect could be ruled out. For protein identification and quantification, DIA analysis against the library generated a quantitative proteome containing 6,195 proteins (**Table S4**) from these tissues in total, averagely 4,835 proteins per sample (**Figure 2B**). Among all the quantified proteins, 62% were based on 3 unique peptides, while the default maximum unique peptides used for protein quantification in Spectronaut are just set at 3 (**Figure 2C**). These data made a solid base for further qualitative and quantitative analysis.

### Proteomics characteristics of gastric cancers

To get a glimpse of the overall pattern from all the samples, the filtration of proteomic data was conducted through criteria described in **Methods**, and were resulted in 4,945 proteins (**Table S4**). These protein abundances in the individual samples were compressed to 96 two-dimensional data points via t-SNE and visualized by scatter plot (**Figure 3A**). The figure revealed that the t-SNE derived distances seemed not to distinguish different subtypes of gastric cancer according to the overall protein abundance patterns, meanwhile, tumors and adjacent tissues presented clearly different patterns in protein abundance.

**Figure 3.**
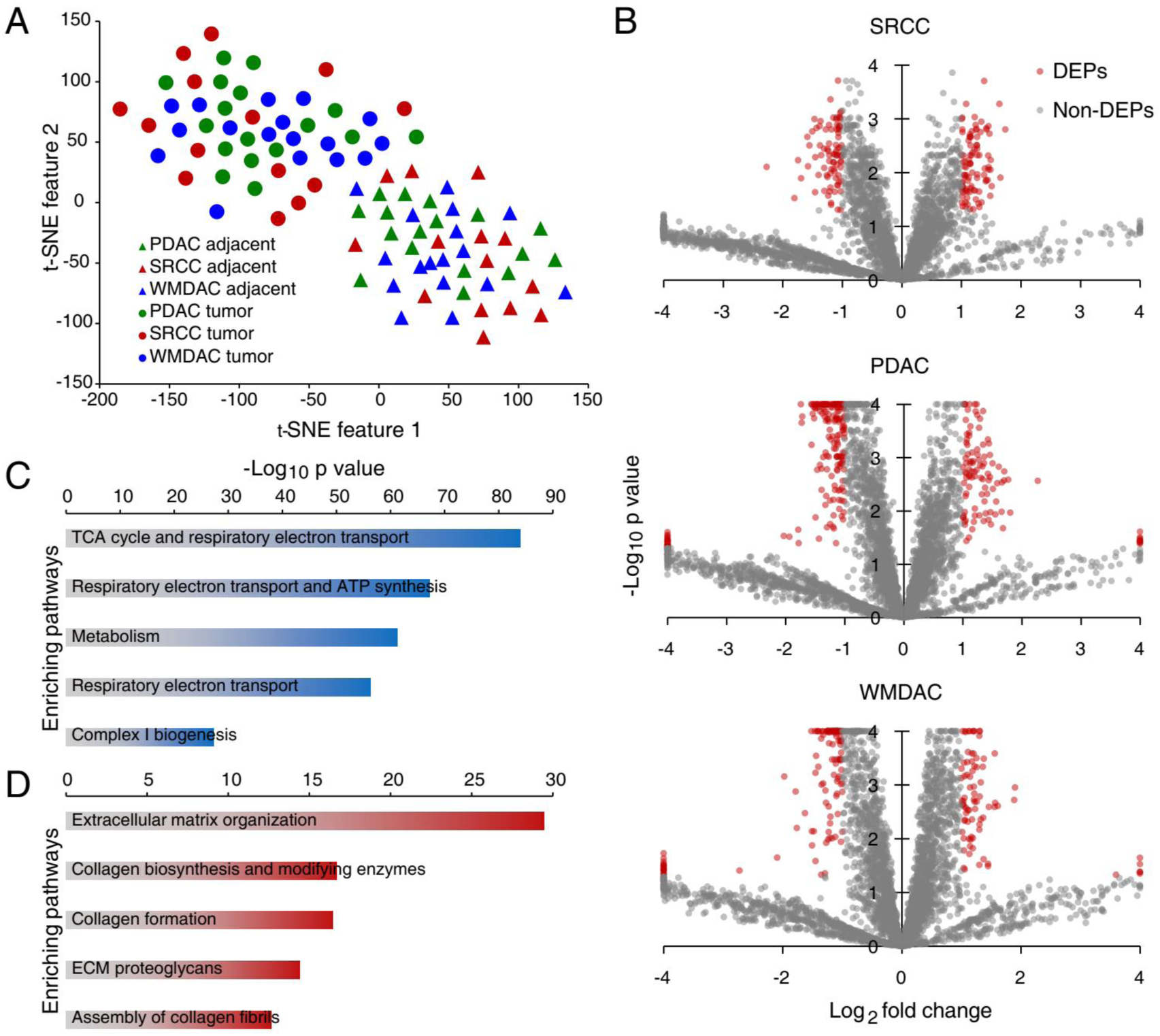
Basic information of the quantitative proteomics in the gastric tissues. A) T-SNE analysis towards the protein abundance gained from the tumor and adjacent tissues in the 3 GC subtypes. B) The presence of T/A-DEPs in 3 GC subtypes on volcano plots based on protein abundance changes and t test. C) and D) The enriched pathways at top 5 for the down regulated and up regulated proteins in all the GC tissues.

Based on the criteria and cutoffs described in **Methods**, we were able to identify the T/A-DEPs, 574/263, 530/235 and 468/213 (down-regulated/up-regulated) from SRCC, PDAC and WMDAC, respectively (**Figure 3B** and **Table S4**). The overlap status of T/A-DEPs were assessed in **Figure S5**, showing 30.9% (380) T/A-DEPs shared by the 3 subtypes. In query of the functions related to the 380 common T/A-DEPs, pathway enrichment analysis was performed using the Reactome pathway database (**Figure 3C** and **D**, **Table S5**). On the bases of evaluation by FDR-adjusted p values produced by the Fisher’s exact tests, extracellular matrix (ECM) organization, collagen biosynthesis and modifying enzymes as well as collagen formation were the top 3 pathways that were commonly upregulated in all the subtypes. As for the enriched pathways with the down-regulated T/A-DEPs, TCA cycle, respiratory electron transport and metabolism were the 3 most pronounced ones. It was a common phenomenon that activation of ECM modification and suppression of aerobic metabolism were well recognized hallmark behaviors of many tumors^14, 15^. This result implied that gastric cancers had major and common differences between their tumors and adjacent tissues which pointed to disruptions of ECM and energy metabolism.

### Comparison of the proteomic characteristics among SRCC and ACs

We further inquired to whether there was any subtype-based abundance feature. In an attempt to hierarchically cluster the 48 cases based on their tumor/adjacent protein ratios (**Figure S6**), it was not easy to distinguish individual subtypes from each other. Then the protein abundances were compared in linear regression and the closeness was evaluated by Pearson correlation coefficient (R^2^). As shown in **Figure 4A**, PDAC and WMDAC were mutually more similar in protein fold change pattern, R^2^ = 0.79, as compared with PDAC to SRCC, R^2^ = 0.70, and WMDAC to SRCC, R^2^ = 0.66, implying that the proteomic abundance of PDAC and WMDAC was generally comparable, whereas that of SRCC was unique to some extent. Based on this overview, more investigations were conducted to pinpoint the proteins as well as pathways with different expression patterns between SRCC and ACs.

**Figure 4.**
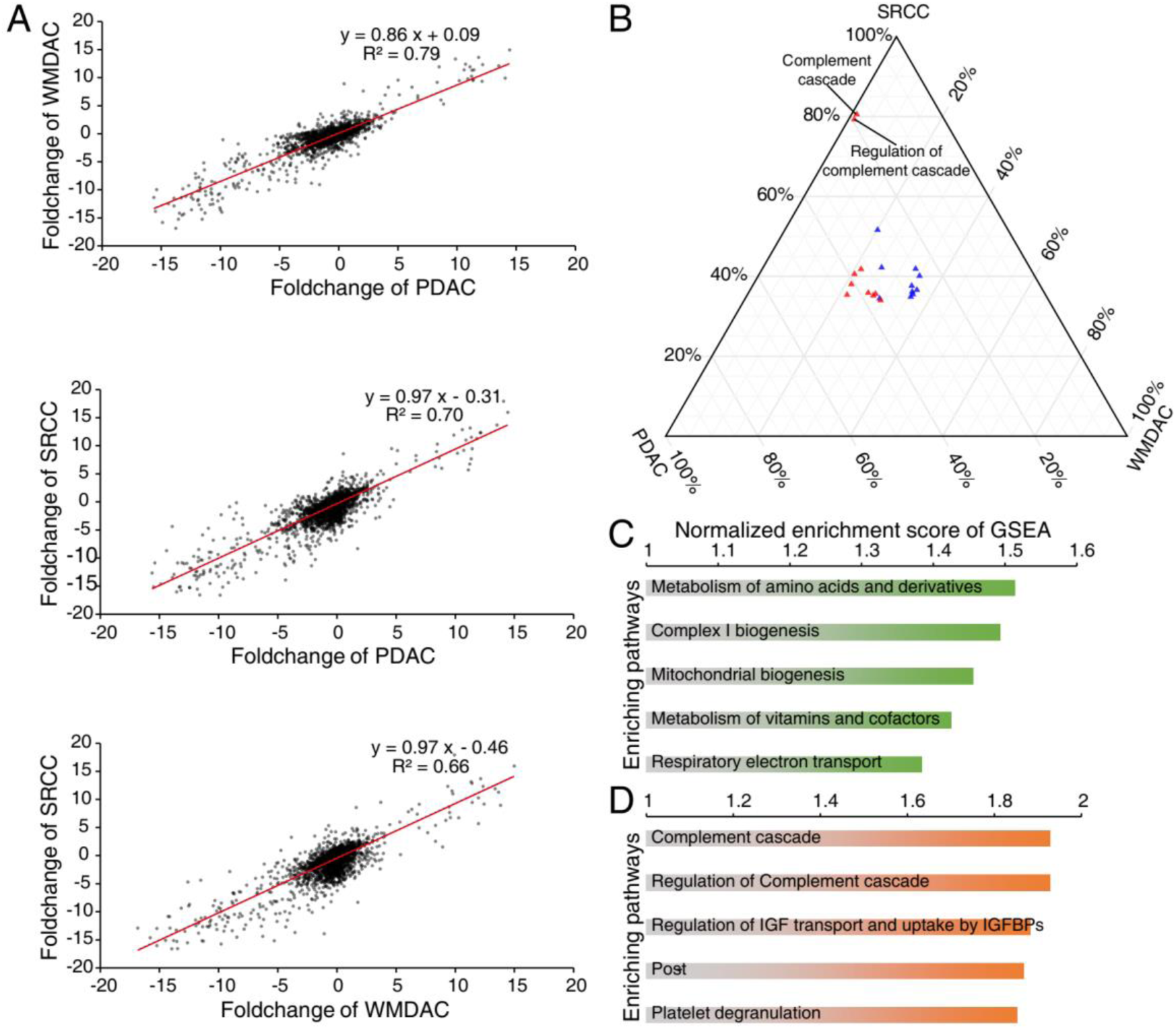
The proteomic characterization of SRCC. A) Linear correlations of the protein abundance ratios (T/A) inter-subtypes. B) Ternary plot indicating subtype specificities of top 10 pathways enriching upregulated T/A-DEPs (red) and downregulated T/A-DEPs (blue). C) and D) The AC-specific and the SRCC-specific pathways at top 5 based on GSEA.

According to the definition of S/A-DEP, of 4,133 candidate proteins, only 10 proteins matched with the criteria (**Table S6**), 6 proteins with higher abundances in SRCC, carcinoembryonic antigen-related cell adhesion molecule 5, matrix Gla protein, mucin-2, mucin-5B, ribosomal RNA processing protein 1 homolog B and serpin B6, while 4 proteins with higher abundances in AC, cytochrome c oxidase assembly protein COX11, mitochondrial 28S ribosomal protein S11, mitochondrial peptidyl-tRNA hydrolase 2 and selenoprotein H (**Figure S7**). In order to find pathways whose regulations were different among subtypes, we carried out pathway enrichment on T/A-DEPs identified from 3 subtypes (**Table S7**). The log10 FDR-adjusted p values of the top enriching pathways were normalized and plotted to ternary scale (**Figure 4B**), demonstrating that the complement cascade and its regulation pathway were specifically upregulated in SRCC. Furthermore, GSEA was used to mine pathways harbored proteomic signals distinguishing SRCC from AC (**Table S7**). The top AC-specific pathways were mostly mitochondrial functions related. While SRCC-specific pathways were majorly related to extracellular reactions including the complement cascade (**Figure 4C and D**), which agreed with previous analysis. The complement cascade involves 138 proteins according to the Reactome database, of which approximately 60% (78) were quantified in the gastric tissues (**Figure 5**). As over one third of the gastric complement proteins exhibited higher abundance in SRCC and the average abundance ratios of T/A for complement proteins were 2 folds more than that in PDAC and WMDAC (**Figure S8**), we came to a deduction that the proteins involved in complement cascade were largely regulated in the SRCC microenvironment, while such observation in the study of gastric cancer was not reported yet. To conclude, despite the overall similar pattern observed among the 3 subtypes, handful proteins were found to express differentially between SRCC and ACs. Meanwhile, pathway enrichment results consolidated that complement cascade was much more upregulated in SRCC than AC.

**Figure 5.**
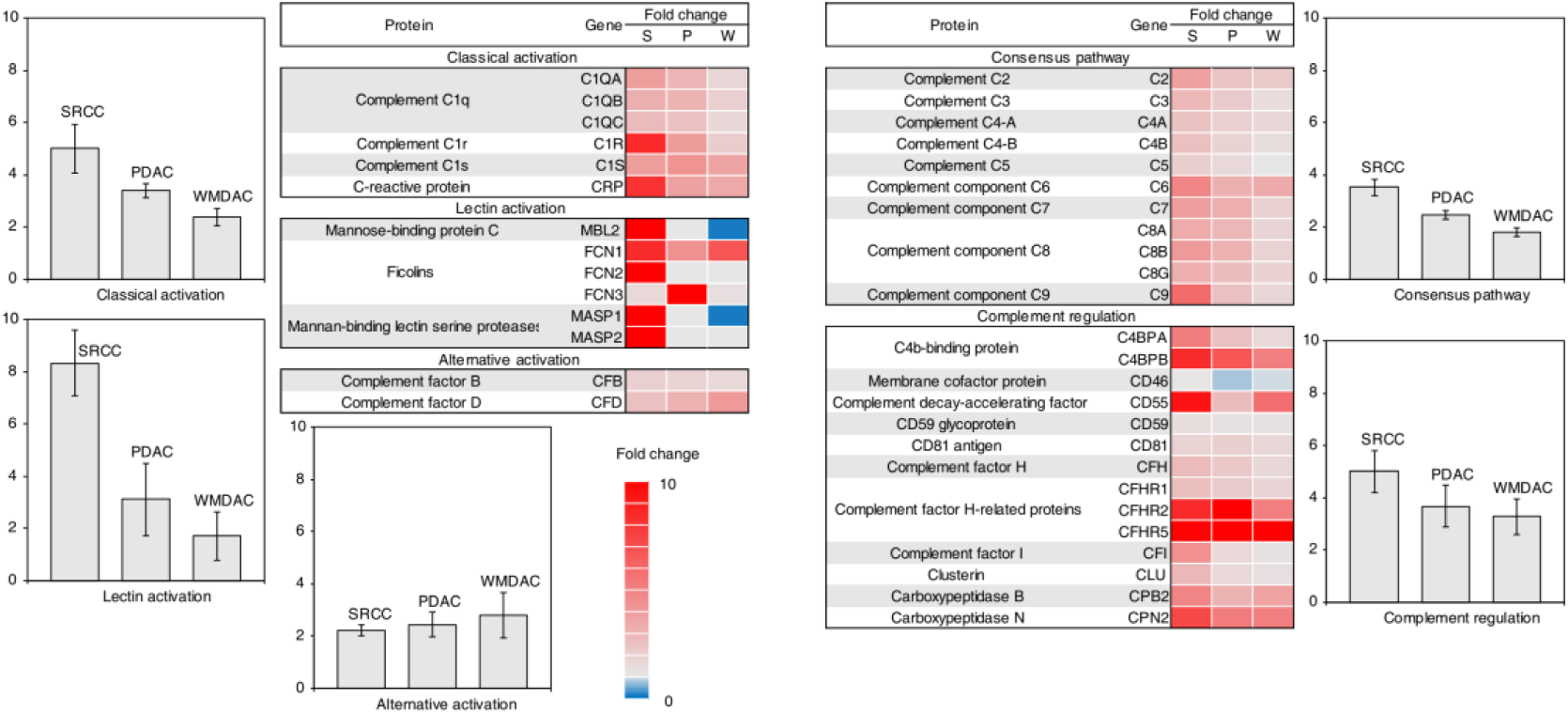
Expressional levels of complement cascade in 3 subtypes. Complement related proteins quantified in 3 subtypes of gastric cancer were grouped into 5 segments, namely, classical activation, lectin activation, alternative activation, consensus pathways and complement regulation. Protein fold changes were indicated by heatmap and average fold change for each segment was described in corresponding bar plot.

### The complement relevant proteome events in SRCC

The complement cascade is harbored in human innate immunity, which likely consists of two events in cancer, complement activation followed by consensus amplification in tumor microenvironment and complemental regulation proteins (CRPs) function in membrane bound or secreted forms in tumor cells. The complement activation generally takes three distinct pathways, namely classical, lectin and alternative, while all the activated pathways finally merge into consensus amplification to exert the cascaded influence of innate immunity. As shown in **Figure 5**, the bulk of proteins in classical and lectin pathway were identified in SRCC with significant upregulation but not in AC tissues, except FCN3, whereas only two proteins of alternative pathways were perceived in all the tissues of SRCC and ACs with insignificant changes of their protein abundance. Moreover, a large amount of immunoglobins that might recognize the tumor-specific antigens and bind to C1 complexes in classical pathway were identified with increased abundance (**Table S4**), implying the activation of classic pathway in SRCC. Mucins (MUC2 and MUC5B) that are the secreted glycoproteins with rich N-acetylglucosamine moiety^16^ and are liganded with lectins^17, 18^ detected in high abundance were significantly upregulated in SRCC as compared that in ACs, implicating that lectin pathway was indeed activated in SRCC (**Figure 5**). In consensus amplification, complement proteins were upregulated to higher extent in SRCC than ACs, whereas the protein fold changes in the pathway appeared less than that in classical and lectin activation (averagely 5.00 fold increase in classical, 8.32 in lectin and 3.51in consensus). All the proteomic evidence thus led to a deduction that the classical and lectin pathways but not alternative pathway were activated in SRCC. The activation signals should be enlarged through consensus amplification, however the changes of protein abundance in consensus pathway were not fully coordinated with the complement activation. This suggested that the delivery of the activation signals were possibly attenuated in SRCC.

Tumor derived CRPs identified in this study, either membrane bound or secreted, generally function as negative regulators to block the complement cascade. As depicted in **Figure 5**, many CRPs exhibited higher abundance in the SRCC tissues. These upregulated CRPs exhibited two characterizations. First, membrane cofactor protein (CD46), complement decay-accelerating factor (DAF, CD55) and CD59 are the common membrane bound CRPs related with tumor to inhibit complement cascade. Although the three proteins were identified in the gastric tissues, only CD55 was found abundance increased in SRCC, but not the others. Second, over 10 secreted CRPs were identified with abundance augment in SRCC. For instance, there were C4b-binding protein (C4bp) and complement factor I that bind or cleave C3/C5 convertases^19, 20^, complement factor H and its related proteins (FHRs) that target and degrade C5 convertase^21^ and carboxypeptidase N and clusterin that inactivate the membrane attach complex (MAC)^22, 23^. Importantly, these secreted CRPs showed significant higher abundance in SRCC against the corresponding adjacent tissues, while their fold changes in SRCC were obviously larger than that in ACs, C4bp (6.75/3.95), DAF (9.28/4.22), factor I (4.39/1.41), clusterin (2.87/1.42) and carboxypeptidases N (6.03/4.22), respectively. Hence, the proteomics evidence supported the postulation that the secreted CRPs in the tumor cells of SRCC were greatly expressed and secreted to matrix, which might mainly response to complement activation in cancer microenvironment and effectively attenuate the pathway of complement consensus amplification.

A question is naturally raised how the complement activation coordinates with complement regulation because of both events with the enhanced abundance of the involvement proteins. We hypothesize a molecular scenario that in tumor microenvironment the complement activation, like classic and lectin, are triggered by degradation products of phagocytosis, chemiotaxis of inflammatory cells or tumor cell lysis. Once the complement activation components are deposited on tumor cell surface, the defense systems within them would be stimulated to exhibit complement-avoidance, by either DAF or a set of the secreted CRPs. Therefore, in SRCC tissues a balance between complement activation and regulation is remained so that some tumor cells escape from complement mediated cytotoxicity (**Figure 6**).

**Figure 6.**
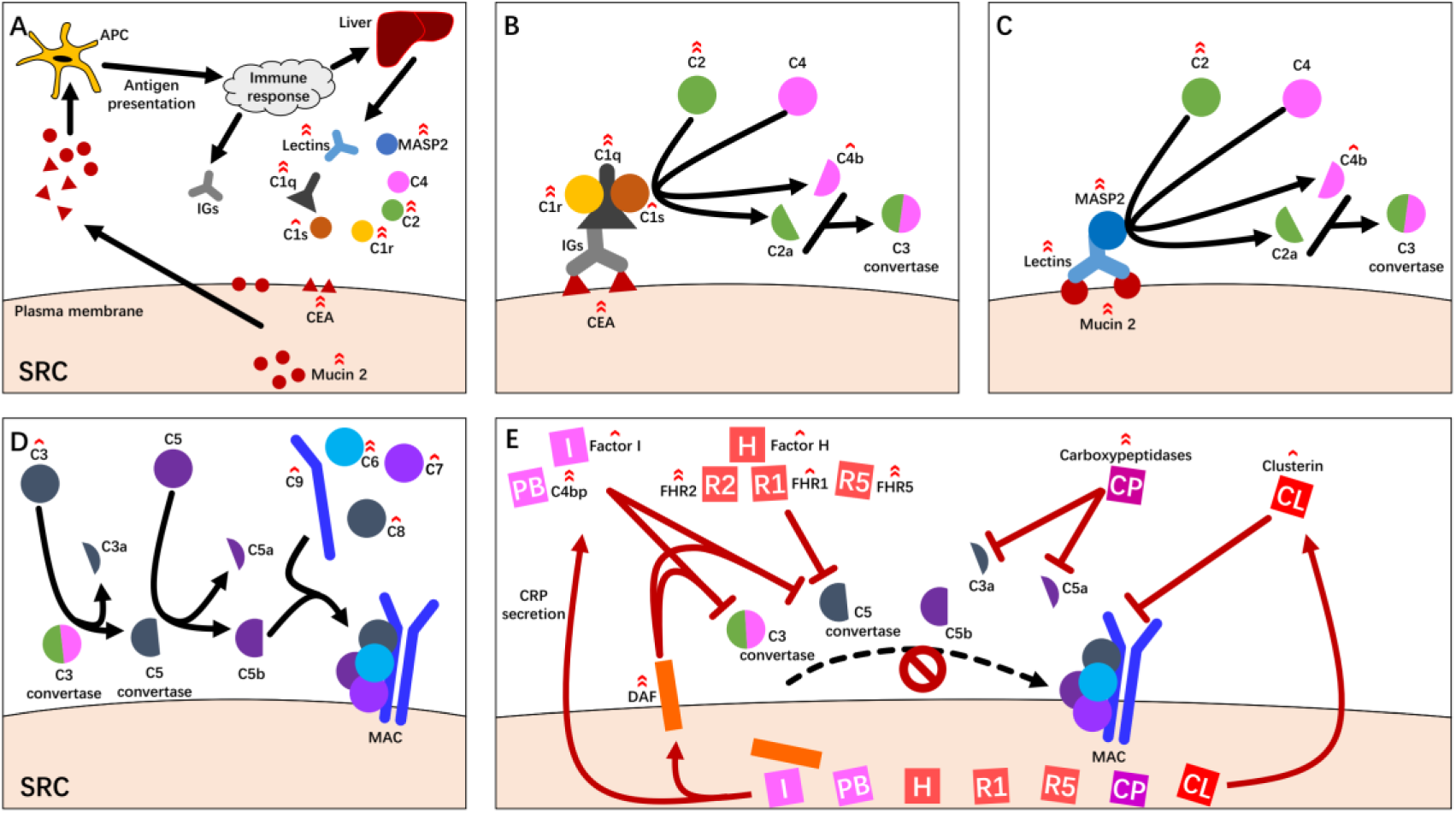
Postulated complement molecular events occurred in SRCC context. Red arrow marks indicate upregulation of corresponding molecules, with single mark indicating fold change > 2 but ≤ 4, and double marks indicating fold change > 4. **A**. Pre-complement reactions. The signet ring cells (SRC) overexpress mucin 2 and CEA, which are captured by antigen presenting cells (APC) in the SRCC microenvironment. The dendritic cells present those markers and activate immune response, which produce IGs to target SRC, and cytokines to stimulate hepatocytes which in turn express complement molecules including lectins, MASP2, C1q, C1s, C1r, C2 to C9 (not completely depicted). **B**. Classical activation. IGs bind to SRC antigen like CEA and recruit C1q, C1r and C1s to form C1 complexes which cleave C2 and C4. The products of this reaction form C3 convertase (C2aC4b complex). **C**. Lectin activation. Complement related lectins bind to SRC surface glycoproteins like Mucin 2 and recruit MASP2 which functions the same as C1 complex described in **B**. **D**. Consensus pathway. C3 convertase cleave C3 into C3a and C5 convertase (C3b). C5 convertase cleave C5 into C5a and C5b. C5b recruit C6 to C9 and form MAC. **E**. Complement regulation. To survive the complement cascade induced cytotoxicity, SRC express DAF, Factor 1 and C4bp to inhibit C3 and C5 convertase, Factor H, FHR1, FHR2 and FHR5 to inhibit C5 convertase, Carboxypeptidases to inhibit C3a and C5a, and Clusterin to abolish MAC.

### Comparison of the proteomic characteristics between gastric cancer tissues and cell lines

Many cell lines derived from the tissues of gastric cancer are widely accepted in academic investigation. After cell proliferation in many generations and the special treatment of cell immortalization, a question has remained whether those cell lines still keep the molecular characteristics of gastric cancer. In this study we tried to seek the answer through comparison of proteomic characteristics between tissues and cell lines. A total of 6,639 proteins were quantified (**Table S8 and Figure S9B**) from all the cell lines, and on average 5,213 proteins were perceived in an individual cell line (**Figure S9A**). The globally normalized protein abundance data were hierarchically clustered with an unsupervised mode as shown in **Figure 7A**, demonstrating no obvious hierarchical cluster because over 50% of the proteins had relatively comparable abundance, whereas the other proteins possessed diverse distribution of their abundance.

**Figure 7.**
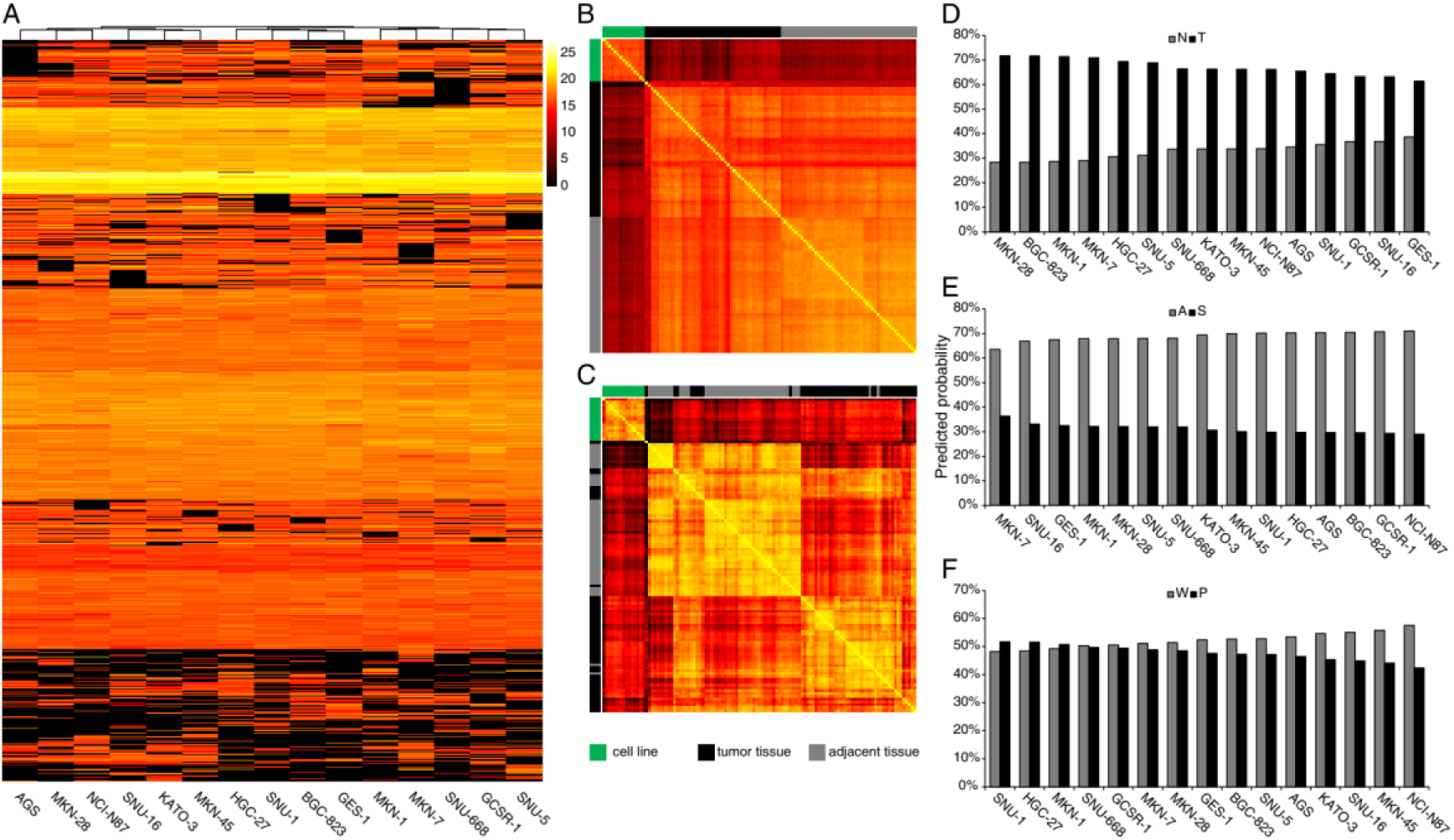
Analysis of the quantitative proteomes in 15 human gastric cell lines. A) Clustering of the quantified proteins in all the cell lines. B) and C) Parallel abundance comparison of cell lines and tissues via Jacard index and correlation coefficient. D), E) and F) Similarity predictions towards GC cell lines and tissues by NT, AS and PW classifier.

The comparability assessment towards proteomic data was carried out in both qualitative and quantitative information. For qualitative comparison, Jacard index^24^ was gained by the ratio of the overlapped proteins to the total proteins in any two samples, and resulted in a Jacard matrix. As illustrated in **Figure 7B**, the Jacard index mean (0.64) for proteins between tissue and cell samples were much less than the values of 0.82 or 0.82 for the proteins within tissues or within cell lines, suggesting that the overall features in the tissue proteome was incomparable with that in cell lines. There were 1,409 proteins uniquely identified in cell lines and 965 uniquely in tissues (**Figure S10A**). Through the pathway enrichment analysis, the unique proteins in cell lines were significantly concentrated in 86 Reactome pathways and those in tissues were enriched into 53 Reactome pathways (**Table S9**), whereas the converged pathways in tissues were completed different from cell lines, strongly endorsing the conclusion drawn from **Figure 7B** and **Figure S10B**. For quantitative comparison, a correlation matrix of protein abundance (**Figure 7C**) was generated from correlation coefficients (R^2^) of the co-identified proteins between tissues and cell lines. Similar to the results of Jacard matrix, the mean R^2^ of 0.57 between tissues and cell lines was obviously smaller than the mean R^2^ within tissues (0.81) or cell lines (0.80), implying the quantification distribution of proteomes was largely different between tissues and cell lines.

Machine learning classifiers are efficient means to find out similar or dissimilar groups in large data. Among a variety of algorithms, random forest^25^ classifier is able to smartly weight and combine the intrinsic input features, thus generalizing reasonable predictions. There were 3 random forest classifiers that were constructed, 1) NT classifier was trained on data from all the 96 LCM samples to classify a cell line into “normal” or “tumor”, 2) AS classifier was trained on data from 48 tumor LCM samples to classify a cell line into “SRCC” or “AC” and 3) PW classifier was trained on data from 34 adenocarcinoma LCM samples to classify a cell line into “PDAC” or “WMDAC”. Cross validations were carried out to find that all 3 classifiers yield acceptable accuracy with the whole dataset (**Figure S11A** and **B**, **Table S10**). Probability of 50% was set as the threshold for class prediction. As a result, all the cell lines of gastric cancer selected were classified to “tumor” with probabilities of 62%∼73% by the NT classifier (**Figure 7E**), and 3 best representatives for tumor were MKN-28, BGC-823 and MKN-1 with probabilities over 71%. Although the GES-1 cell line derived normal gastric epithelia, it was also classified into “tumor” due to its predicted probability was 62%, GES-1 had the lowest probabilities to be “tumor” in all the cell lines, implying that it was still different from tumor tissue somehow. All the cell lines were classified to “adenocarcinoma” with probabilities of 63%∼71% by the AS classifier with the top 3 representatives of AC, NCI-N87, GCSR-1 and BGC-823 (**Figure 7F**). Surprisingly, 3 cell lines derived from SRCC tumors, KATO-3, GCSR-1 and SNU-668 (**Table S1**) were also recognized as AC. As for PW, the prediction probabilities generated by this classifier were ranged from 48%∼58% (**Figure 7G**), which were too close to 50% to reach an acceptable predictions, suggesting that the cell lines for PDAC or WMDAC were not well grouped through PW classifier. Based upon these classifiers, we came a conclusion that the 14 selected cell lines of gastric cancer appeared similar proteomic features with the AC in tissues, nevertheless the cells currently used for SRCC study were incomparable with the correspondent tumor tissues based upon proteomic features at least.

## Discussion

Previously, in term of the depth, the best result obtained from MS based proteomics on gastric cancer was reported by Ge et al^26^, who managed to quantify ∼4,400 proteins per sample on average and ∼9,200 proteins for a total of 168 samples. This was done by feeding large amount of peptide samples and fractions (∼100 μg in 6 fractions) to LC-MS/MS with DDA. In comparison, we applied DIA strategy in this study to quantify ∼4,800 proteins per sample on average and ∼6,100 proteins for a total of 96 samples, achieving slightly better quantification results per sample yet but much improved inter sample comparability. We noticed that, in spite of the similar scale and depth achieved by the present work and Ge’s work, there are differences in the DEPs-enriching pathways concluded by the two works. To investigate, we compared the pathways enriching DEPs from the 3 histological subtypes in our work and 3 molecular subtypes, PX1, PX2 and PX3 classified by Ge et al. As listed in **Table S11**, the upregulated pathways of 3 subtypes in our work were mainly ECM related, and the downregulated pathways were mainly energy metabolism related. In contrast, the PX1 subtype didn’t clearly imply its downregulated pathway, meanwhile the PX2 and PX3 showed the downregulation pathways related with energy metabolism and translation, respectively. As for the upregulated pathways, the PX1 and PX2 concentrated in transcription and cell cycle related functions, while the PX3 exhibited enrichment of immune systems related pathways. The inter study differences in DEPs and their enriching pathways could be attributed to two factors, 1) different schemes of subtype classifications adopted by the two works may highlight different functional aspects for each subtypes and 2) LCM used our work reduced interfering signals from other types of cells present in tumors, while Ge’s work made use of bulk tissues for the analysis. Nevertheless, it should be recognized that the results of both studies reflected only parts of the gastric cancers and further investigations featured with advanced scale and depth are needed to fully characterize the gastric cancer.

With an emphasis on the SRCC’s unique characteristics in comparison to ACs, 10 proteins were revealed to have distinct expression patterns between SRCC and ACs in this work. To examine whether these patterns were supported at transcription level, a transcriptomic gastric tumor dataset including 407 samples (32 normal tissues, 363 AC tumors and 12 SRCC tumors) generated in a TCGA project (TCGA-STAD, https://portal.gdc.cancer.gov/projects/TCGA-STAD) was retrieved. The mRNA abundances in FPKM were normalized and values for the 10 relevant genes were extracted, shown in **Figure S14**. Among them, CEACAM5, MUC2, MUC5B and MRPS11 had similar SRCC/AC differences in their transcripts and proteins, which made them solid SRCC specific indicators. Carcinoembryonic antigen-related cell adhesion molecules, encoded by CEACAM5 gene, had long been recognized as a tumor associated transmembrane protein. Its overexpression was observed in gastric and colon cancers^27, 28^. Besides its intercellular adhesive role played in various types of tissues^29^, CEACAM5 also possess a series of tumor promoting functions such as disruption of cell polarization, inhibition of cellular differentiation and anoikis^30-32^. Biomarker study carried out by Zhou et al. associated CEACAM5 expression with worse prognosis of gastric cancer^33^. When it comes to SRCC, the presence of CEACAM5 was not consistent. Immune staining results demonstrated in Terada’s study suggested CEACAM5 had higher level of expression in gastric and colorectal SRCC^34^, while Warner et al. reviewed 20 prostate SRCC cases only to find 4 CEACAM5 positive cases^35^. Nevertheless, as this study and independent TCGA dataset revealed that CEACAM5 was specifically highly expressed in gastric SRCC comparing to AC, there is a potential opportunity to develop unique therapy to SRCC by targeting CEACAM5 whose protein product is located at tumor cell surface. In fact, such strategy was already conceptualized and experimented for colorectal cancer^36^. Mucin 2 and mucin 5B are 2 mucus comprising proteins widely produced and secreted by epithelial goblet cells under physiological condition. One of the functions of mucin 2 is suppression of inflammation occurs at mucous epithelia, deficit of which was postulated to be a promoting factor of colon cancer^37, 38^. However, in the case of gastric SRCC, the overexpression of the secreted mucins doesn’t necessarily contribute to positive effect, since a significant amount of mucins are stored in the intracellular droplets of signet ring cells, which potentially indicates a disruption of physiological secretion of mucins. Further examination of expression levels of the mucin secretion related proteins, including rab3 GTPase-activating protein, protein unc-13 homologs, protein unc-18 homologs, syntaxins, synaptotagmins, synaptosomal-associated proteins and vesicle-associated membrane proteins^39^ in our proteomics data, didn’t support this postulation. The unique morphology of SRCC complicates the function of overexpressed mucins, and one can hope future investigations harness the complication in treating SRCC. The mitochondrial ribosomal small subunit 11, encoded by MRPS11, was shown to be expressed at a specifically lower level in SRCC in comparison to AC. The expression of MRPS11 was correlated with favored outcome in colorectal cancer^40^, but its functional association with cancer is yet to be discover.

As emphasized in the results, complement cascade and its regulation were found to be characteristically upregulated pathways (**Figure 6**). Concerning the cancer related complement cascade deregulation as reviewed by Afshar-Kharghan^41^, many previous studies has been carried out, covering glioblastoma, melanoma as well as cervical, ovarian, lung, colorectal, breast and thyroid bladder. The complement cascade carried out double-sided functions in development of various tumors. On the one hand, it promotes elimination of tumor cells by activating adaptive immune systems and forming MAC in microenvironment which directly induces apoptosis in tumor cells, on the other hand, the complement cascade promotes proliferation of tumor cells via anaphylatoxin signaling. For complement cascade in gastric cancers, very limited findings were available. Chen et al revealed that expression of complement proteins C5b, C6, C7, C8 and C9 was tumor-related and differentiating stage-dependent in gastric adenocarcinoma^42^, while Inoue et al reported that a complement regulator, CD55 was constantly expressed higher in gastric cancer cells than the normal gastric tissues^43^. Other complement cascade related proteins, C1r, C1s, C3 and the most central C4b, as well as multiple complement regulators lacked documentations until the present study. As regards complement-related proteins in gastric SRCC, only C1q was reported to be associated with the tumor development^44^. For the first time, our study discovered a bulk of the proteins in complement cascade pathways highly sensitive in SRCC tissues. Although the implications of complement cascade are incomplete and naive, its importance in host immune system has attracted studies related to a wide range of diseases. As pointed out by Kleczko et al^45^, therapies targeting complement cascade had already been experimented against immune system related diseases like rheumatoid arthritis and age-related macular degeneration. Judged by the quantitative proteomes profiled by the present study, it was assumed that the evading of complement induced cell death by up-tuning the complement negative regulators was an significant characteristic of SRCC, and targeting the CRPs might be an effective approach to inhibit SRCC.

Although we discovered the association of complement cascade activation to gastric SRCC in a subtype-constrained manner, it should be noted that the proteins involved in complement cascade were largely missing in any of the 14 gastric cancer cells. In fact, about 80% (61) of proteins in complement cascade in gastric tumor tissues were not reflected by any gastric cancer cell lines. This caveat needs to be aware when cell lines are used to model tumors, where molecular events occur in the tumor microenvironments such as complement cascade, are lost in cell lines.

## Acknowledgments

This work was supported by the funding from the National Key R&D Program of China (2017YFC0908403) and the National Key Basic Research Program of China (973 program) (No. 2014CBA02002 and No. 2014CBA02005).

