## Supplemental information for "Protein profiling reveals the characteristic changes of complement cascade pathway in the tissues of gastric signet ring cell carcinoma"

**Supplemental information 1** Detailed methods

Sample collection

Frozen tissues were obtained from the specimen library of digestive diseases in Xijing Hospital, China. All the tissue samples in the specimen library were frozen in liquid nitrogen at -196℃ immediately after surgical resection. According to pathological records, 2,522 cases of gastric tumors samples were stored between 2011 and 2016. Considering the relative low incidence of SRCC, we adopted a strategy to first select the SRCC cases followed by inclusion of PDAC/WMDAC cases by matching with the SRCC cases in terms of age and gender (**Figure 1)**. The tissues diagnosed as SRCC by pathologists were further confirmed by microscopic recheck with hematoxylin and eosin (H&E) (BA4025, BASO, China) staining. In order to select the matched cases of PDAC and WMDAC, the cases for each subtype were selected in a case-wise matching manner to the SRCC cases as described **Figure 1**. Since the incidences tend to be distributed at lower age in gastric SRCC than ACs, low-age cases with available frozen tissues and > 50% cancer cells for the AC subtypes were supplemented. The inter-subtype matching goodness in terms of age, gender, T and N staging was measured by Wilcox test[1] (age), and $\chi^2$ test[2] (gender, T and N staging). The paired tumor and adjacent tissues for those cases were collected for this study.

The cell lines in **Table S1** were collected from multiple sources, including a gastric epithelia-derived GES-1, and 14 gastric tumor-derived GCSR-1, AGS, NCI-N87, KATO-3, HGC-27, BGC-823, MKN-1, MKN-7, MKN-28, MKN-45, SNU-1, SNU-5, SNU-16 and SNU-668.

LCM

To get the tumor cells isolated from other cells, the tissues were first thawed and embedded in optimal cutting temperature compound (4583, Sakura, USA), then were frozenly sectioned with Cryostat (CM1900, Leica Biosystems, Germany) at -20 ℃, resulting in 8 μm-thick tissue sections. The adjacent tissues in a sectioning sequence were evenly divided and mounted to PEN membrane glass slides (LCM0522, Applied Biosystems, USA) and were polysine-coated glass slides (6776215, Thermo Scientific, USA), as target slides and indicator slides, respectively. The tissue sections on slides were slightly fixed with 75% ethanol for 1 minute, and were stored at -80 ℃ until use. The indicator slides were H&E stained to guide LCM of their target slides, while LCM were operated on the target slides to isolate tumor cells corresponding to their sample subtypes, as well as epithelial cells for the adjacent tissue (ArcturusXT LCM system, Applied Biosystems, USA).

Peptides preparation

Lysis buffer composed of 2% RapiGest (186001861, Waters, USA), 100 mM $NH_4HCO_3$ (A610032-0500, BBI LIFE SCIENCES, China), 2 mM EDTA (E0105-500g, Sangon Biotech, China) and 1 mM PMSF (P0754-25G, Sangon Biotech, China) was used for lysis of LCM tissues and protein dissolution, followed by ultrasonication (VCX130, Sonics & Materials, USA). DTT (0281-BEJ-100G, AMRESCO, USA) at 20mM and IAM (I6125-10G, SIGMA, USA) at 55 mM were sequentially introduced for reduction and alkylation of the protein disulfide bonds. The dissolved proteins were digested with 2 μg trypsin (V5280, Promega, USA) in 100mM $NH_4HCO_3$ at 37 °C overnight. The resulted peptides were desalted with self-packed tip columns with OLIGO R3 reverse-phase resin (1133903, Applied Biosystems, USA). Since the DIA analysis depends on DDA data derived spectral library, one tenth of each tumor and adjacent peptide sample was aliquoted and pooled for generation of DDA. The pooled peptides were fractionated to 10 fractions with self-packed tip columns using high-pH Xtimate C18 resin (01710-01100, Welch, China). The peptides were quantitated with Pierce quantitative fluorometric peptide assay kit (23290, Thermo Scientific, USA). The peptides in the cell lines were processed through the similar preparation procedure.

LC-MS/MS

Approcimately1.5 μg peptides spiked in 15 ng iRT peptides (Ki-3002-1, BIOGNOSIS, Switzerland) in each sample were loaded to an Ultimate 3000 nanoLC system (Thermo, USA), which was equipped with a self-packed column (30 cm x 150 μm with 1.8 μm C18 resin). An 120 min elution gradient (**Figure S1**) was run followed by a nanoESI interface to introduce peptides to an Orbitrap Fusion Lumos mass spectrometer (Thermo, USA). Selection range for precursor ions was set 400 – 1500 m/z and HCD at 30% relative energy. For DDA runs, precursor ions were scanned with orbitrap analyzer at 60,000 resolution then top 30 most intensive ions were selected for fragmentation. The fragmented ions were scanned with orbitrap analyzer at 15,000 resolution. AGC was set to targets of 1e5 (MS) and 2e4 (MS/MS) with max accumulation time of 50 ms. For DIA runs, precursor ions in sequential windows of 25 m/z were fragmented and scanned in orbitrap analyzer at 30,000 resolution. AGC was set to target of 5e4 and max accumulation time 54 ms. DIA samples were analyzed with duplicates, and in random order to avoid potential batch effect.

Data processing

Before database search, the post translation modifications (PTMs) of the data were surveyed by Preview V3.0 (PROTEIN METRICS). According to the result (Table S2), Carbamidomethylation artifacts (+57 on N-terminus, H and K), Deamidation (+1 on N and Q) and DTT addition (+152 on C) were used as variable modifications in following database search. The DDA data were searched against Swiss-Prot human proteome (20,381 entries) by DDA search engine operated in Spectronaut Pulsar X 12.0 (BIOGNOSIS, Switzerland) with setting at most 3 missed cleavage(s) for tryptic peptides, and at most 5 top variable modifications. The DIA data were treated with DIA search engine against the ion library self-built in the same software following the parameters recommended by the vendor.

The protein quantification data for all runs were globally normalized to an equal total, i.e. the mean sum of all runs. Then the normalized data for each of the three subtypes, SRCC, PDAC and WMDAC, were strictly filtered and their differentially expressed proteins (DEPs) between tumor and adjacent tissue (T/A-DEPs) were identified through procedure depicted by **Figure S2**. The data for each of the three subtypes, SRCC, PDAC and WMDAC, were filtered and their differentially expressed proteins between tumor and adjacent tissue (T/A-DEPs) were identified through procedure depicted by **Figure S2**. In brief, each protein in each case (tumor / adjacent pair) were classified according to its abundances in corresponding tumor and adjacent samples. Proteins with enough (more than 2/3 of all cases) full data points (cases with available data from tumor and adjacent samples, class 1 in **Figure S2**), or enough tendency-agreed unilateral cases (cases with data from either tumor or adjacent samples, class 2 and 3 in **Figure S2**) plus cases whose absolute $\log_2$ transferred ratio > 1 (class 1.1 and 1.2 in **Figure S2**), were retained while other proteins were filtered out. A protein with enough full data points was determined as T/A-DEP if its FDR-adjusted p value of t test (calculated with build-in functions in R 3.4.4) < 0.05 and absolute $\log_2$ transferred fold-change > 1, and all the proteins with enough tendency-agreed unilateral cases plus cases whose absolute $\log_2$ transferred ratio > 1 were designated as T/A-DEPs. The pathway enrichment analysis on T/A-DEPs were achieved with g:Profile[3] on Reactome[4] pathways through over representation analysis (FDR adjusted p value < 0.05).

The proteins that passed the filtration as above and presented in all subtypes were firstly selected, then the abundance ratios for all the selected proteins were attained, in SRCC termed RS and in AC as RA. To determine the DEPs between SRCC and AC (S/A-DEPs), 2 comparisons were conducted, comparison of protein abundance (SRCC/AC), and comparison of abundance ratios (RS/RA) between SRCC and AC. An S/A-DEP was defined once its FDR-adjusted Wilcox test p value < 0.05 and its absolute $\log_2$ transferred fold-change > 1 in both comparisons.

The differential pathways between SRCC and AC were achieved through Gene Set Enrichment Analysis (GSEA)[5] based on the protein abundance ratios of tumor/adjacent.

Aiming to classify cell lines into gastric cancer subtype, a random forest model was trained upon the protein abundance in tissues (**Figure S3**), in which the proteins (square root n) ranked at abundance were randomly picked up 1,000 times for training the classifier using scikit-learn module[6], python 3.7. data in were used by machine learning which. Cross-validation in “leave one out” manner was used to seek the features subset with highest accuracy. Protein abundance data from cell lines were treated in the same manner, then were fed to the trained classifier for their tissue-type likelihood prediction.

Other software for data analysis

The data were reshaped and accommodated for analyzing purpose with Excel 2016 (Microsoft, USA), R 3.4.4, Pandas module[7] and NumPy[8] module of Python 3.7. The data of t-distributed stochastic neighbor embedding (tSNE)[9] were generated by scikit-learn module of Python. The heatmaps were generated and visualize by pheatmap package of R, while the other data were plotted by Excel 2016 and GraphPad Prism 8.11.

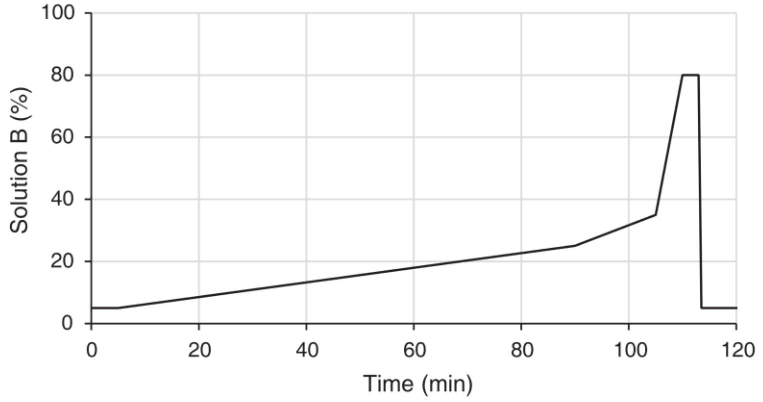


| Time | Phase B (%) |
|---|---|
| 0 | 5 |
| 5 | 5 |
| 90 | 25 |
| 105 | 35 |
| 110 | 80 |
| 113 | 80 |
| 113.5 | 5 |
| 120 | 5 |

**Figure S1** LC gradient used in LC-MS experiments. Binary phase was applied, with phase A being 0.1% FA+2% ACN and phase B being 0.1% FA+98% ACN.

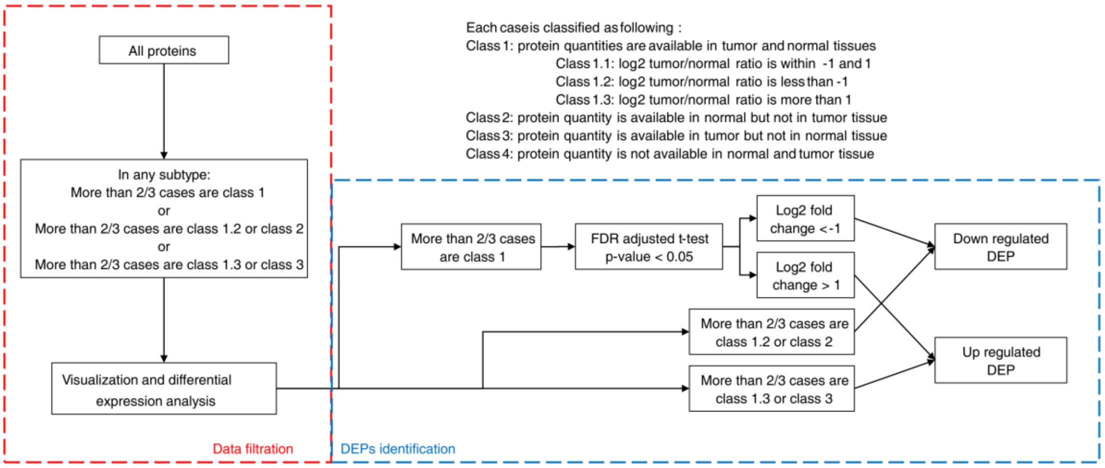


**Figure S2** Filtration of data and definition of T/A-DEPs.

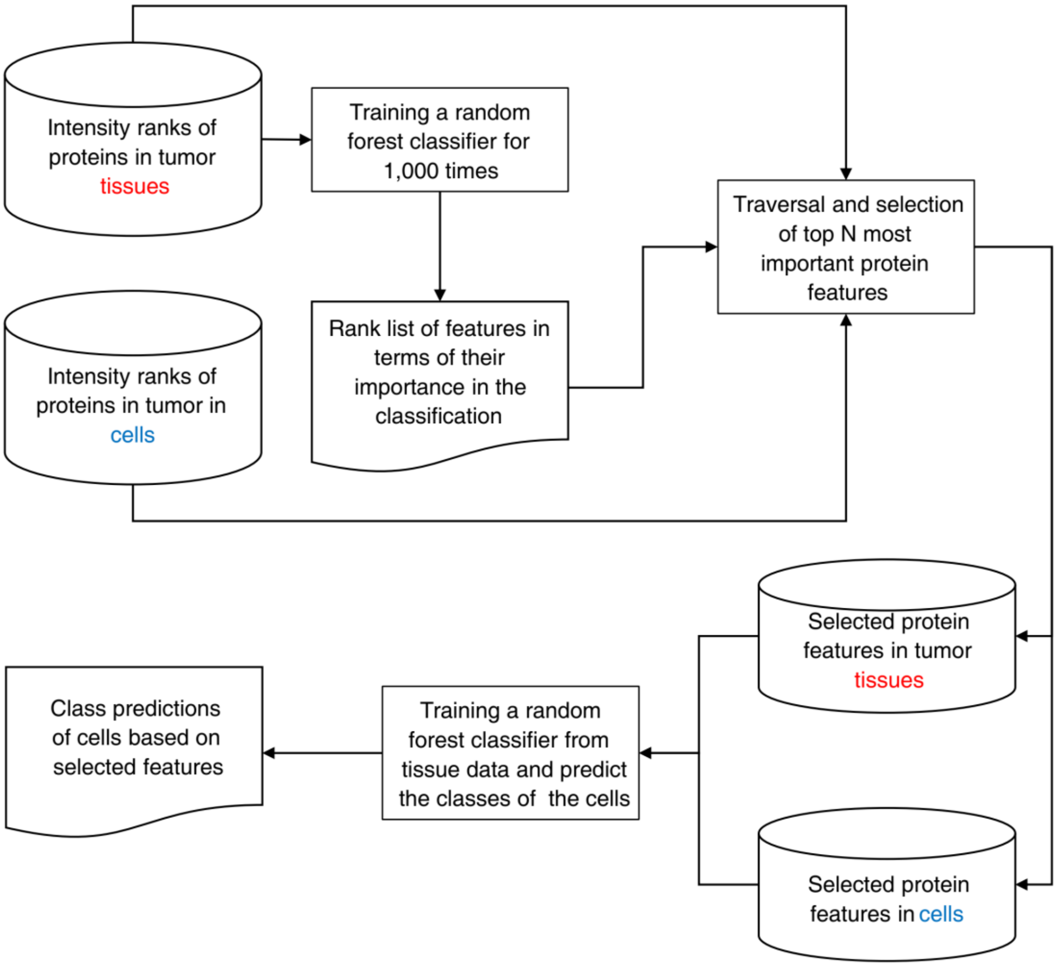


**Figure S3** Flow chart of training, cross validating and applying machine learning (random forest) classifier.

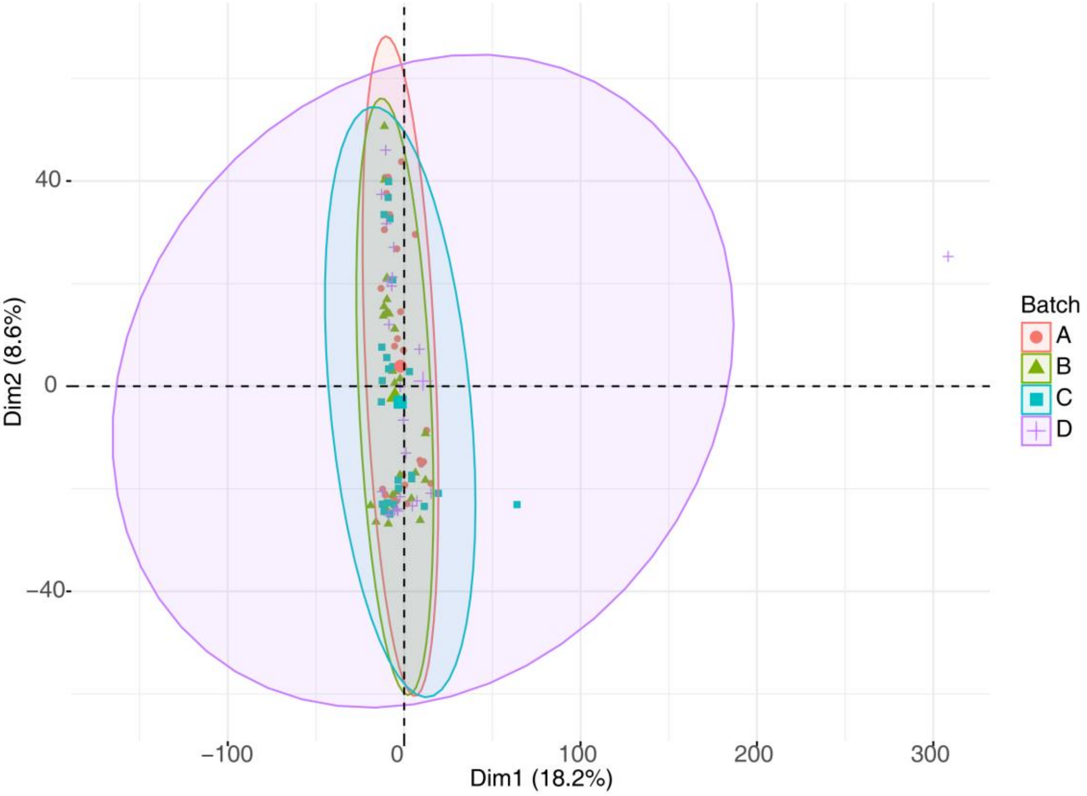


**Figure S4** PCA visualization of the whole dataset grouped with batches. Ellipses indicate confidence intervals.

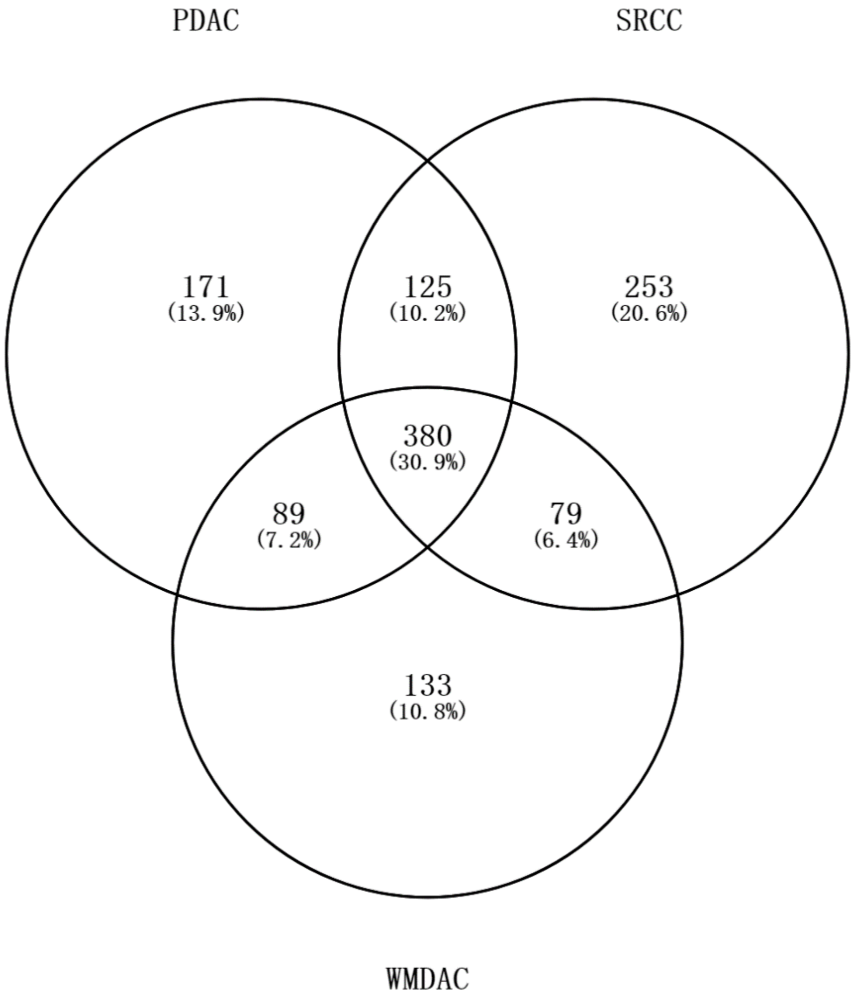


**Figure S5** Venn chart of T/A-DEPs discovered in 3 subtypes of gastric cancer.

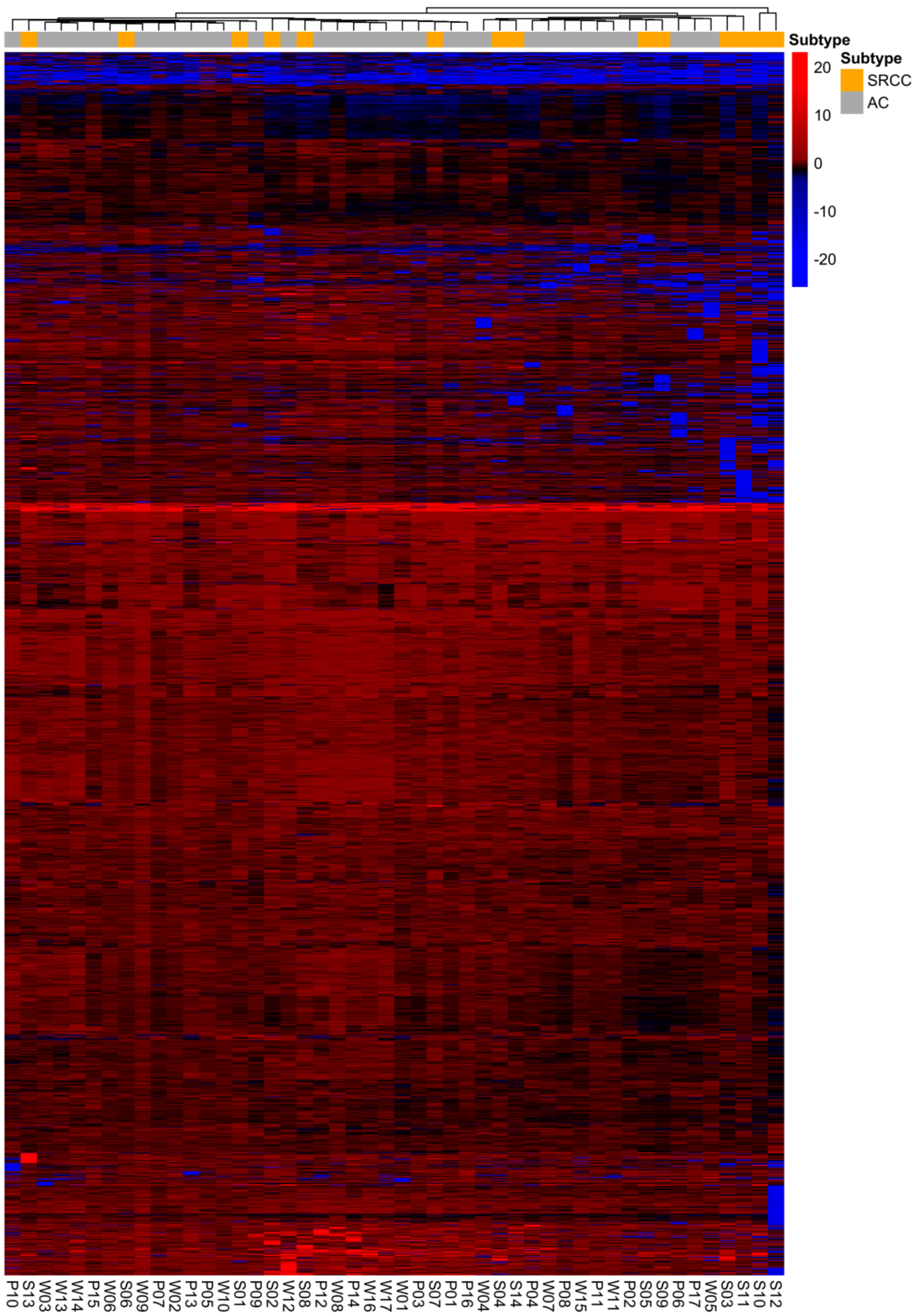


**Figure S6** Heatmap for T/A ratios. Columns are samples and rows are proteins passed the filtration. Samples are labeled by SRCC and AC. Columns and rows are hierarchically clustered.

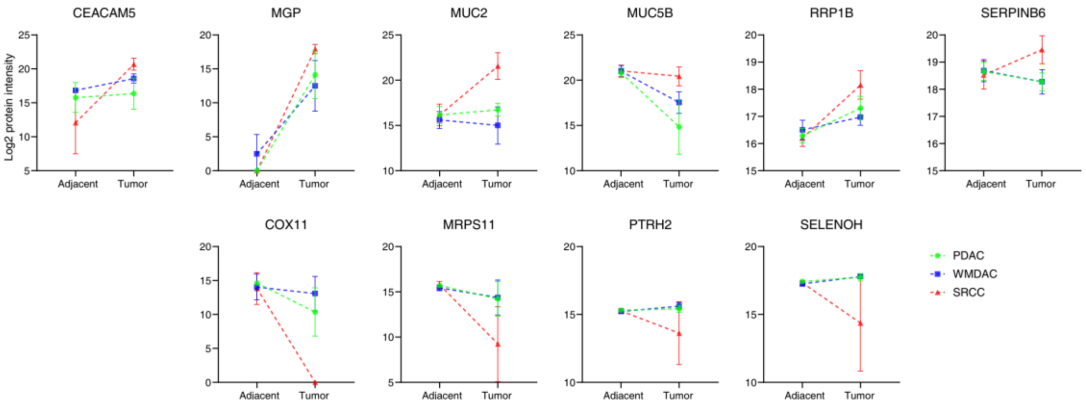


**Figure S7** Expression statuses of 10 S/A-DEPs. In each line chart, x axis is tissue type and y axis is $log_2$ transformed protein abundance. Data are grouped by 3 gastric cancer subtypes. Error bars indicate confidence intervals.

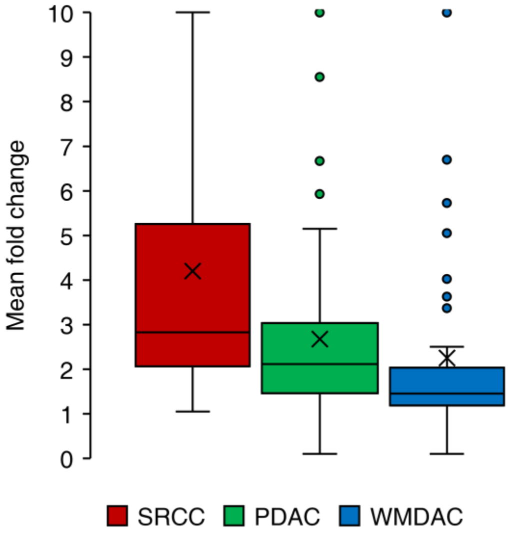


**Figure S8** Overall expression levels of complement cascade in 3 subtypes of gastric cancer. Cross marks indicate the mean values.

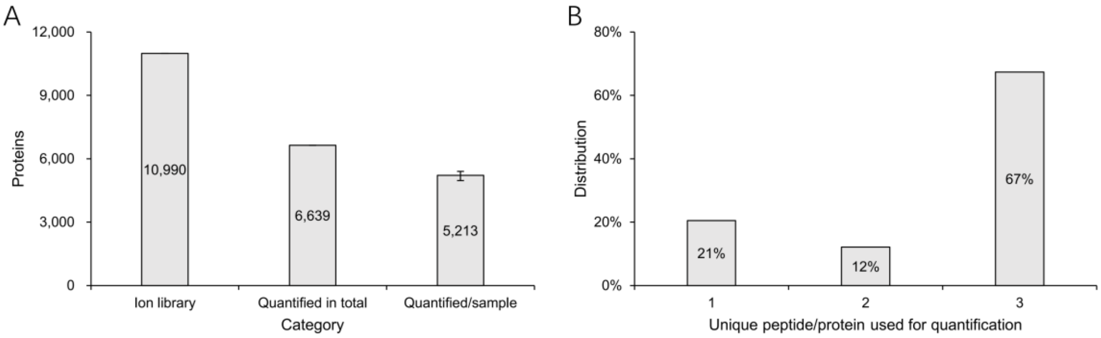


**Figure S9** Data quality of cell lines. B) The proteins in the gastric cell lines identified using DDA and DIA approach (error bar indicates the upper and lower bound of proteins quantified/sample). C) Distribution of the unique peptides in the quantified proteins.

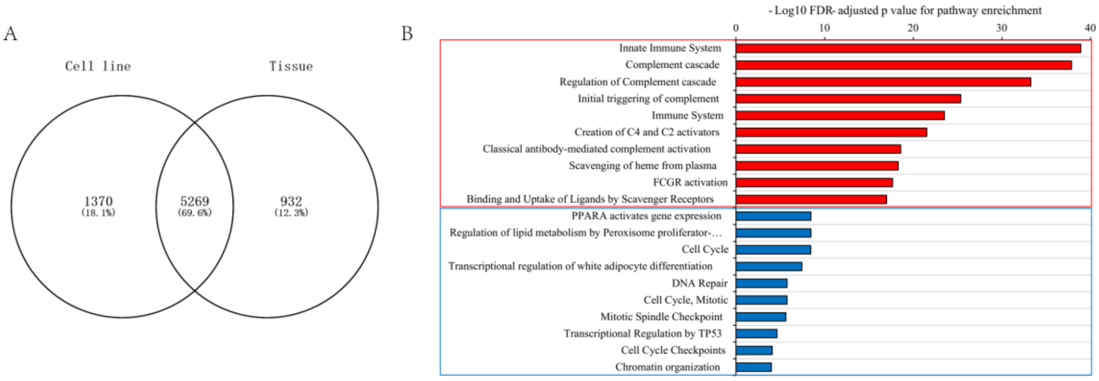


**Figure S10** Differences between gastric tissues and cell lines regarding quantified proteins and pathways. A) Venn chart of quantified proteins in gastric tissues and cell lines. B) Pathways enriching unique proteins in gastric tissues and cell lines in A. Top 10 pathways were listed. Red: Tissue unique, blue: cell line unique.

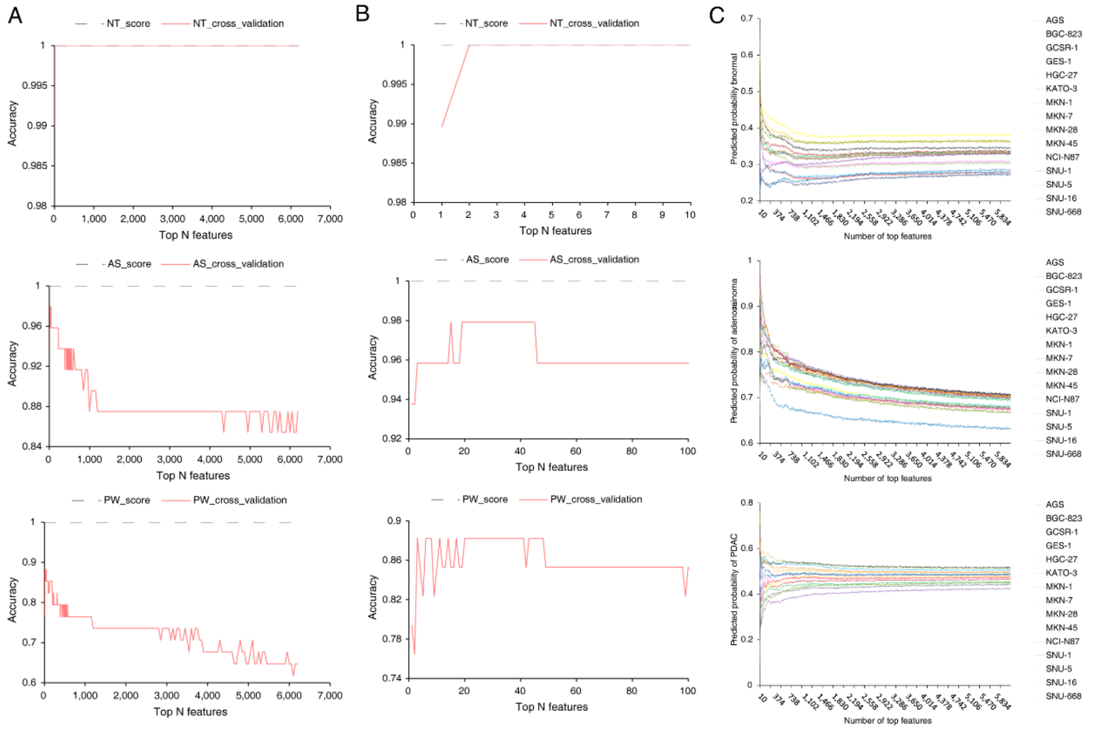


**Figure S11** Effects of using different sizes top features on the random forest classifiers. A) and B), cross validation derived accuracies in whole and zoomed scales, respectively. C) Predicted probabilities. Upper, middle and lower figures are NT, AS and PW classifiers.

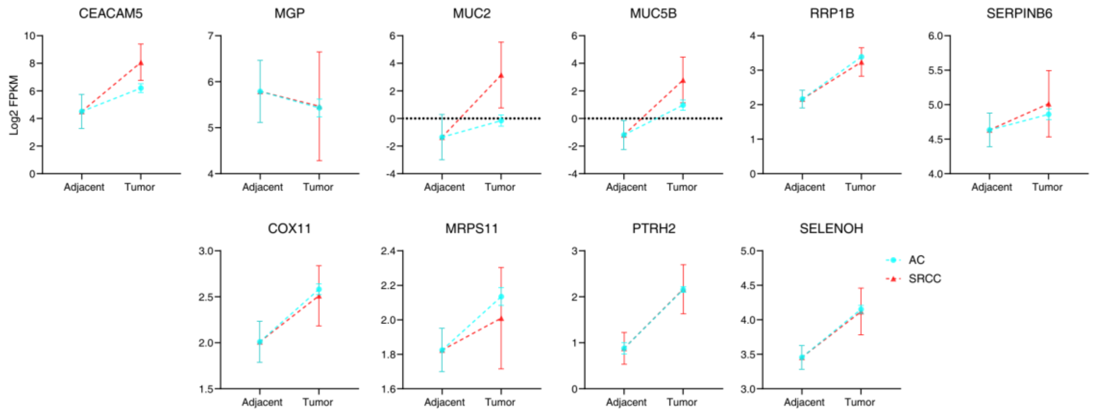


**Figure S12** Expression statuses of 10 S/A-DEPs in TCGA dataset. In each line chart, x axis is tissue type and y axis is $\log_2$ transformed FPKM. Data are grouped by 3 gastric cancer subtypes. Error bars indicate confidence intervals.
